# Genome Assembly of *Docynia delavayi* Provides New Insights into α-Farnesene Biosynthesis

**DOI:** 10.1101/2024.05.06.592716

**Authors:** Jinhong Tian, Zhuo Chen, Can Jiang, Siguang Li, Xinhua Yun, Chengzhong He, Dawei Wang

**Author notes:** Correspondence author at: Key Laboratory for Forest Resources Conservation and Utilization in the Southwest Mountains of China Ministry of Education, Southwest Forestry University, Kunming, 650224, China. Corresponding (Y.X.H), (H.C.Z); (W.D.W). These authors contributed equally to this work (Jinhong Tian, Zhuo Chen).

## Abstract

*Docynia delavayi* is an economically significant fruit species with a high market potential due to the special aroma of its fruit. Here, a 653.34 Mb high-quality genome of *D. delavayi* was first reported, of which 93.8% of the sequences were assembled to 17 chromosomes. A total of 48325 protein-coding genes were annotated in *D. delavayi* genome. Ks analysis proved that two whole genome duplication (WGD) events occurred in *D. delavayi*, resulting in the expansion of genes associated with terpene biosynthesis, which promoted its fruit-specific aroma production. Combined multi-omics analysis, α-farnesene was detected as the most abundant aroma substance emitted by *D. delavayi* fruit during storage, meanwhile a α-farnesene synthase gene and 15 transcription factors (TFs) potentially involved in α-farnesene biosynthesis were identified. Further studies for the regulation network of α-farnesene biosynthesis revealed that DdebHLH, DdeERF1 and DdeMYB could activate the transcription of *DdeAFS*. To our knowledge, it is the first report that MYB TF plays a regulatory role in α-farnesene biosynthesis, which will greatly facilitate future breeding programs for *D. delavayi*.

## Introduction

*Docynia delavayi* (2n=2x=34) is a fruit tree species belonging to the *Docynia* genus of the Rosaceae family, which is native to Southwest China (Peng et al., 2021). It has been extensively processed into varieties of foods and traditional herbs by local ethnic minorities for its great edible and medicinal value (Xia et al., 2022; Zhao et al., 2012). As a natural green food, its fruit containing various macronutrients and micronutrients which provided a wealth of nutrients for humans (Li et al., 2022a). Especially, its mature fruit possesses intense rosy and floral aromas, which makes people delightful (Wang et al., 2023b). Additionally, its fruit are rich in flavonoid bioactive substances, which play effective roles in anti-tumor and hypoglycemic (Deng et al., 2014; Wang et al., 2023a). To increase the efficiency of conservation and utilization of *D. delavayi* germplasm resource, its genetic diversity, plus-tree selection, mechanism of pericarp color variation, metabolome in leaves and fruit were studied in recent years (Chen et al., 2022; Li et al., 2022a; Peng et al., 2021; Wang et al., 2023b; Wang et al., 2022b; Xia et al., 2022; Xu et al., 2023). However, the lack of genome-wide information considerably limits the conduct of its molecular breeding efforts.

The Rosaceae family contains about 3000 species, most of which are fruit and ornamental tree species that bring great economic benefits to humans (Initiative et al., 2013; Wang et al., 2021). According to the fruit trait, it is subdivided into 4 subfamilies: Maloideae, Prunoideae, Rosoideae and Spiraeoideae (Zhang et al., 2012). However, Rosaceae members exhibit significant phenotypic diversity in which the frequent morphological synapomorphies are not easily recognized, so identification based on the molecular level is imminent (Zhang et al., 2012). To date, more than 20 members of the Rosaceae family have accomplished high-quality genome sequencing, greatly advancing our understanding of the Rosaceae plants (Wang et al., 2021). The whole genome sequencing of *D. delavayi* could provide new evidence for the classification of Rosaceae plants, meanwhile also contributing to understanding its evolutionary history and the molecular basis distinct biological characteristics.

The production of aroma in edible fruits is an indicator of measuring consumer preference and fruit quality (Liu et al., 2021a). The aroma substances are also the signals that they communicate with the environment and other organisms for most fruit trees, helping them to spread seeds and defend against natural enemies (Shen et al., 2023). The aroma in plant mainly derives from massive volatile organic compounds (VOCs), including terpenes, esters, alcohols, aldehydes, ketones, etc., of which terpenoids are the most abundant (Huang et al., 2023). α-Farnesene is one of the simplest acyclic sesquiterpene of terpenes family, which was main biosynthesized through Mevalonate (MVA) pathway in cytoplasm, while being mainly affected by the expression of *AFS* gene and regulated by a variety of TFs (Liu et al., 2021b; Wang et al., 2020). α-Farnesene not only imparts aroma to fruits but is also widely used in industrial and agricultural production (Du et al., 2022; Liu et al., 2021b), so exploring the molecular mechanism of their biosynthesis can effectively increase the scientific utilization value of *D. delavayi*.

Here, a high-quality chromosome-scale genome of *D. delavayi* was first obtained using the combination of the next, third-generation and Hi-C scaffolding technology. Then, using the post-harvest *D. delavayi* fruits as materials and combining the reference genome, transcriptome and volatile metabolome analysis, the candidate genes associated with α-farnesene biosynthesis were identified and verified using RT-qPCR, subcellular localization and dual-lucferase. In summary, this work provides precious genetic information for *D. delavayi* study, and has greatly contributed to its genetic improvement and molecular breeding.

## Results

### Genome sequencing, assembly and annotation

*D. delavayi* is one of the important economic fruit trees belonging to the Rosaceae family (Fig 1A, B), which was selected for sequencing and assembly. Based to the K-mer analysis, the *D. delavayi* genome size was evaluated at 615 Mb, with a repeat rate of 23.7%; the heterozygosity rate was 0.54% belonging to the heterozygous genome; the highest peak was 110 corresponding to the average depth of 17-mer (Table S3; Figure S1). After determining the genome size, a total of 43.65 Gb data (∼72×coverage) was generated using oxford nanopore technologies sequencing with the N50 of 32.94 kb (Table S4). A 653.34 Mb genome was assembled with a contig N50 of 16.25 Mb and the longest contig size of 36.65 Mb (Table 1). The results of BUSCO assessment showed that about 96.5% of complete gene elements were found in *D. delavayi* genome with relatively high credibility of the assembly result (Table 1). Lastly, the Hi-C assisted assembly results showed that the 612.98 Mb genomic sequence data (93.8%) were anchored onto 17 putative chromosomes with a scaffold N50 of 35.15 Mb and GC content of 38.47%, which showed high continuity (Table 1; Fig. 1C). The Hi-C heatmap demonstrate that the distinct regions of 17 chromosomes could be clearly separated, of which the minimum size of chromosomes is 28.79 Mb and the maximum size is 48.61 Mb (Table S5; Fig. S2).

**Table 1.** Statistics of the *D. delavayi* genome assembly and assessment

| Feature | Value | Feature | Value |
| --- | --- | --- | --- |
| Contig N90 | 4.18 Mb | Complete | 96.50% |
| Contig N80 | 9.00 Mb | Complete and single-copy | 60.80% |
| Contig N70 | 12.14 Mb | Complete and duplicated | 35.70% |
| Contig N60 | 13.82 Mb | Fragmented | 0.80% |
| Contig N50 | 16.25 Mb | Missing | 2.70% |
| Longest | 36.65 Mb | Super-Scaffold N50 | 35.15 Mb |
| Total Size | 653.34 Mb | Super-Scaffold N90 | 30.98 Mb |
| Total Number ($\geq 1$ kb) | 115 | Total | 612.98 Mb |
|  |  | Anchored rate (%) | 93.80% |

**Fig. 1.**
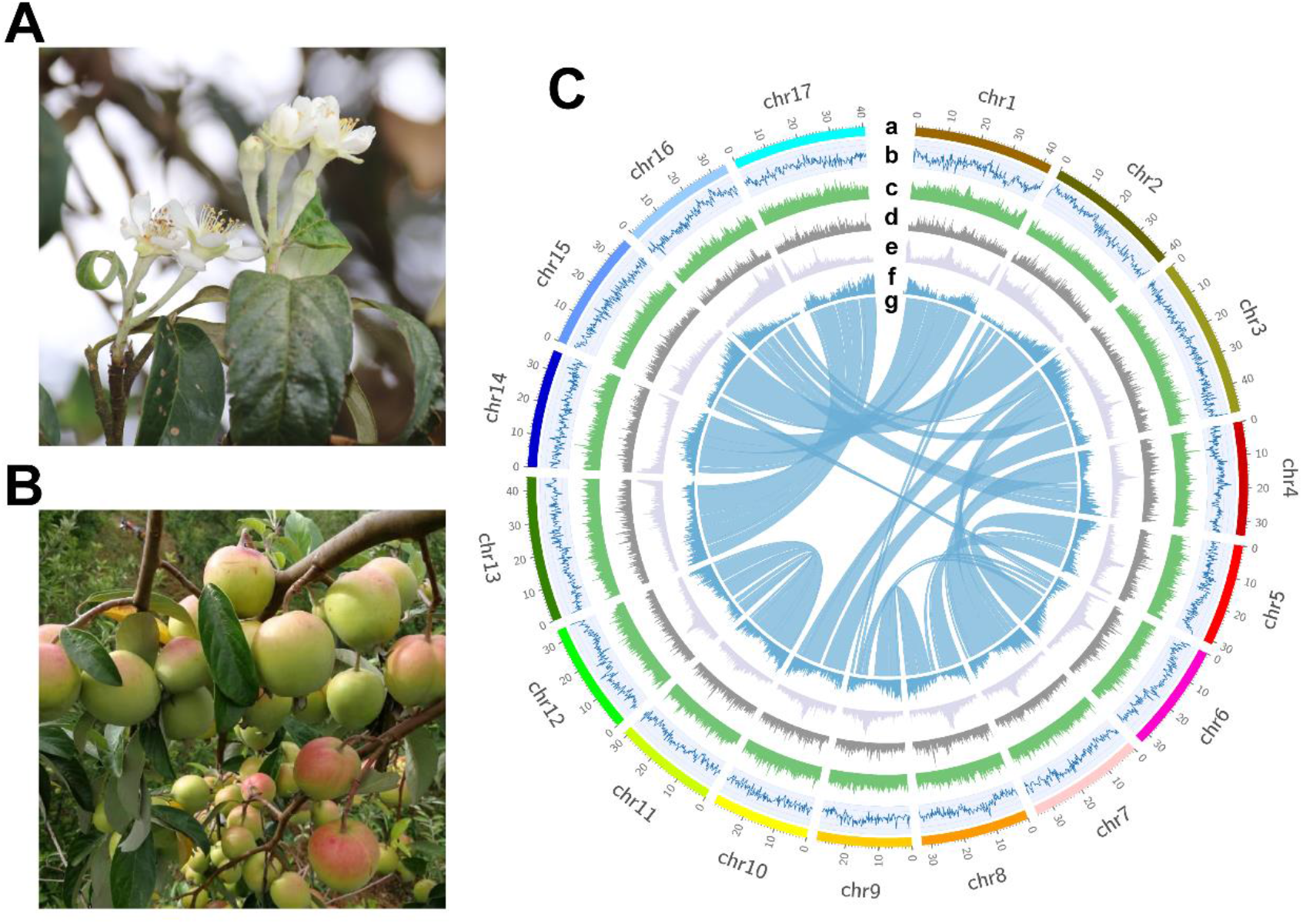
Morphology and the genome overview of *D. delavayi*. A. Flower of *D. delavayi*. B. Fruits of *D. delavayi*. C. a, karyotype analysis; b, GC content; c, protein-coding gene density; d, DNA transposon density; e, LTR/Copia transposon density; f, LTR/Gypsy transposon density; g, synteny blocks in *D. delavayi*.

A total of 347.92 Mb (56.76% of the genome length) repeat regions were identified in *D. delavayi* genome, of which the TR elements accounted for only 3.60% of the whole genome sequence (Table S6). Additionally, LTR retrotransposons (57.27%) was the most abundant type of interspersed repeats, followed by DNA transposon (13.52%), LINE (1.64%) and SINE (0.03%) (Table S7; Fig 1C). In LTR retrotransposons, the Gypsy (30.64%) and Copia (16.33%) were the two most plentiful elements (Fig. 1C). The non-coding RNA were annotated from the *D. delavayi* genome, including 247 miRNAs, 753 tRNAs (transfer RNA), 760 rRNAs (ribosomal RNA) and 686 snRNA (small nuclear RNA) (Table S8). Moreover, a total of 48325 protein-coding genes were predicted from *D. delavayi* genome with an average CDS length of 1041 bp, an average transcript length of 2695 bp, and an average exon number per gene of 4.6. The result of comparative genomic analysis showed that *D. delavayi* has the largest number of protein-coding genes in five Rosaceae plants, while it seemed to be not related with genome size (Table S9; Fig. S3). After gene annotation, the BUSCO evaluation of 1440 highly conserved genes showed that 88.4% genes were identified, reflecting that the prediction quality of gene was high (Table S10). The result of functional annotation showed that 44520 genes out of 48325 genes (95.29%) were successfully annotated via the NR, Swissprot, KEGG, KOG, TrEMBL, Interpro, GO databases (Table S11; Fig. S4).

### Gene family and phylogenomic analysis

A total of 34053 gene families were found in *D. delavayi* genome, of which 2625 were unique to *D. delavayi* (Table S12). Furthermore, a comparison of the number of orthologous genes in 12 species showed that *P. persica* had the least number of multi-copy gene families (6652) and the highest number of single-copy gene families (6068), whereas *M. domestica* contained the greatest number of multi-copy gene families (1761) and the lowest number of single-copy gene families (20146) (Table S13; Fig. 2A). By contrast, *D. delavayi* contained 3619 unique gene families, an intermediate number among the 12 species. The Venn diagram of gene families in the five Rosaceae family species displayed that *R. rugosa, M. domestica, P. communis, P. persica* and *D. delavayi* shared 12383 gene families, whereas 1177 specific gene families were identified in *D. delavayi* (Fig. 2B).

**Fig. 2.**
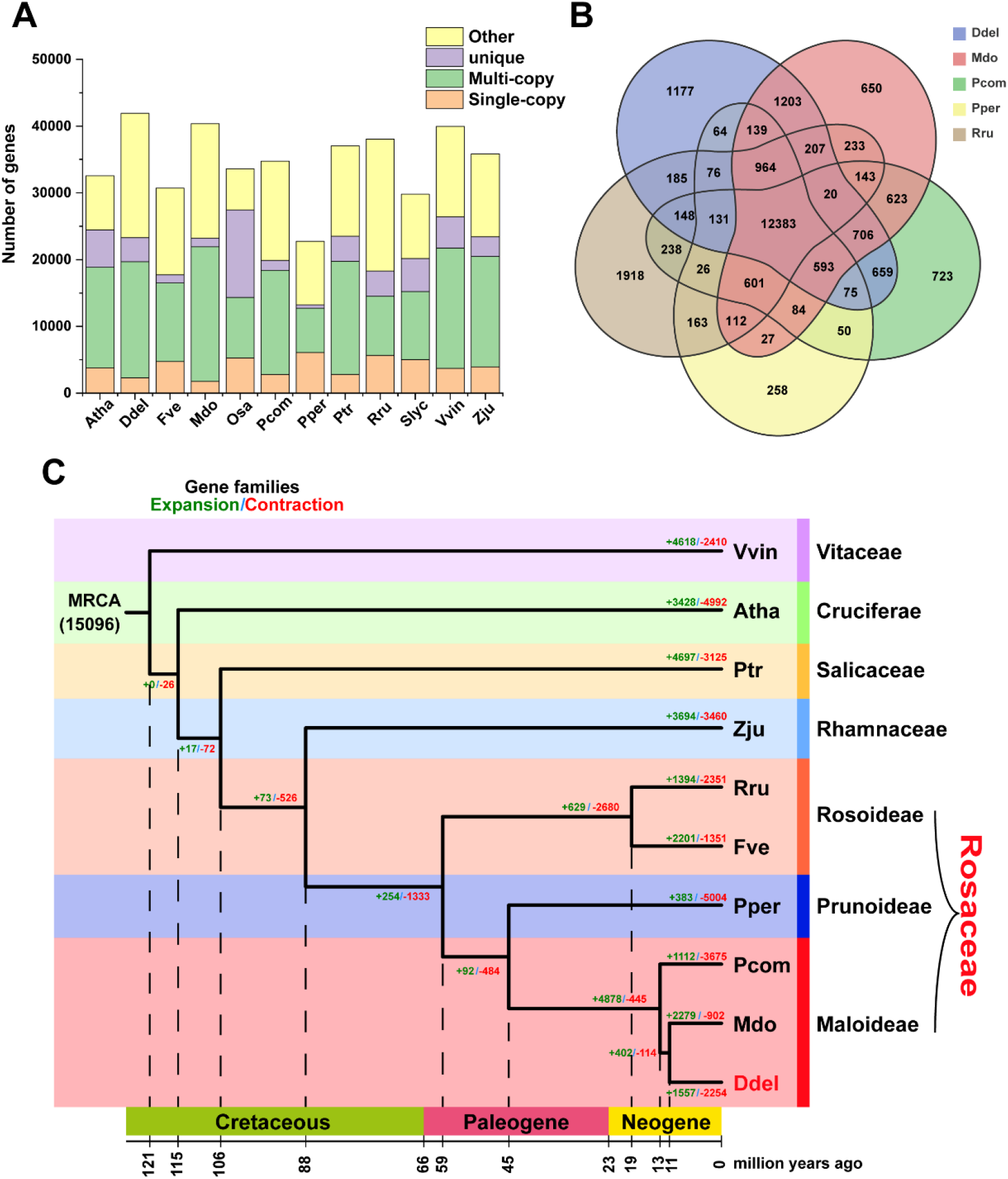
Comparative and evolutionary analysis of the *D. delavayi* genome. A. Number of unique, single-copy and multi-copy genes in 12 representative species. B. Venn diagram of shared orthologous gene families for 5 Rosaceae species. C. Phylogenetic tree, estimation of divergence time and analysis of gene family expansion and contraction of *D. delavayi* genome. The green number represent the number of expansion gene family, and the red number represent the number of contraction gene family. The abbreviations of the *Atha, Ddel, Fve, Mdo, Osa, Pcom, Pper, Ptr, Rru, Slyc, Vvin* and *Zju* represented *Arabidopsis thaliana, Docynia delavayi, Fragaria vesca, Malus domestica, Oryza sativa, Pyrus communis, Prunus persica, Populus trichocarpa, Rosa rugosa, Solanum lycopersicum, Vitis vinifera* and *Ziziphus jujuba*, respectively.

The functional annotation of the unique genes showed that 86 genes were enriched in translation (46), regulation of translation (7), regulation of translational initiation (3), cytoplasmic part (49), cytoplasm (53), intracellular non-membrane-bounded organelle (41), macromolecular complex (58), ribosome (34), ribonucleoprotein complex (37), mannosidase activity (6), mannosyl-oligosaccharide 1,2-alpha-mannosidase activity (4), ribose phosphate diphosphokinase activity (4), structural constituent of ribosome (34) and ribosome binding (5) terms using GO annotation (Fig. S5A), and 87 genes were enriched in ribosome (ko03010, 44), mismatch repair (ko03430, 18), nucleotide excision repair (ko03420, 17), SNARE interactions in vesicular transport (ko04130, 9) and stilbenoid, diarylheptanoid and gingerol biosynthesis (ko00945, 10) pathway by KEGG annotation (Fig. S5B).

The results of phylogenetic tree showed that *D. delavayi* belonged to Maloideae subfamily of Rosaceae family, which was clustered together with *M. domestica*, with an estimated divergence time of 11 million years ago (Fig. 2C). Additionally, Fig. 2C showed that Maloideae and Prunoideae were more closely related than Maloideae and Rosoideae, of which Maloideae diverged from Prunoideae at 45 MYA. Expansion and contraction analysis of 15096 shared gene families from 10 species displayed that 1557 gene families have expanded and 2254 contracted in *D. delavayi* genome. Furthermore, the KEGG annotation of the 1557 expansion gene families showed that a total of 20 pathways were significantly enriched (*P*<0.05), including Terpene backbone biosynthesis (ko00900, *P*<0.01), Monoterpene biosynthesis (ko00902, *P*<0.01) and Sesquiterpene and triterpene biosynthesis (ko00909, *P*<0.01) pathway related to the plant terpene biosynthesis (Fig. S6A). The GO annotation showed that a total of 45 terms were significantly enriched (*P*<0.05) consisting of 19 biological process, 19 molecular function categories and 7 cellular component (Fig. S6B). Notably, 8 genes were annotated as terpene synthase activity (GO: 0010333), suggesting that they probably played important roles in terpene biosynthesis of *D. delavayi* fruit.

### Analysis of genome synteny and WGD

The WGD events during the *D. delavayi* genome evolution was further investigated through the genome synteny and Ks distribution analysis. A total of 493 syntenic blocks containing 16467 collinear gene pairs within *D. delavayi* genome, and we could clearly find large-scale segment duplications between chromosomes, suggesting that it might also undergo the Maloideae-specific WGD event (Fig. 1C). The statistics of gene type displayed that 5625, 2942, 2986, 13554 and 23218 were identified as singleton, proximal, Tandem, dispersed and whole genome duplication genes, respectively (Table S14). The result of the syntenic analysis between *D. delavayi* and *M. domestica, F. vecsa* showed that the *D. delavayi* genome shared 402 syntenic blocks containing 45552 collinear gene pairs with *M. domestica* and 354 syntenic blocks containing 29617 collinear gene pairs with *F. vecsa* (Table S15). In contrast, the syntenic blocks between the *D. delavayi* and *M. domestica* chromosomes had a better fit (Fig. 3A). There were 2 peaks in the Ks distribution of all syntenic gene pairs in *D. delavayi* genome at approximately 0.175 and 1.445, which might correspond to the WGD event shared by the ancestors of dicotyledonous plants and the recent WGD event and, respectively (Fig. 3B). Additionally, the recent WGD event occurred before the differentiation of *M. domestica* and *D. delavayi* but after the differentiation of *D*.*delavayi* and *A. thaliana, F. vesca, P. persica*, which was similar with the phylogenetic analysis result.

**Fig. 3.**
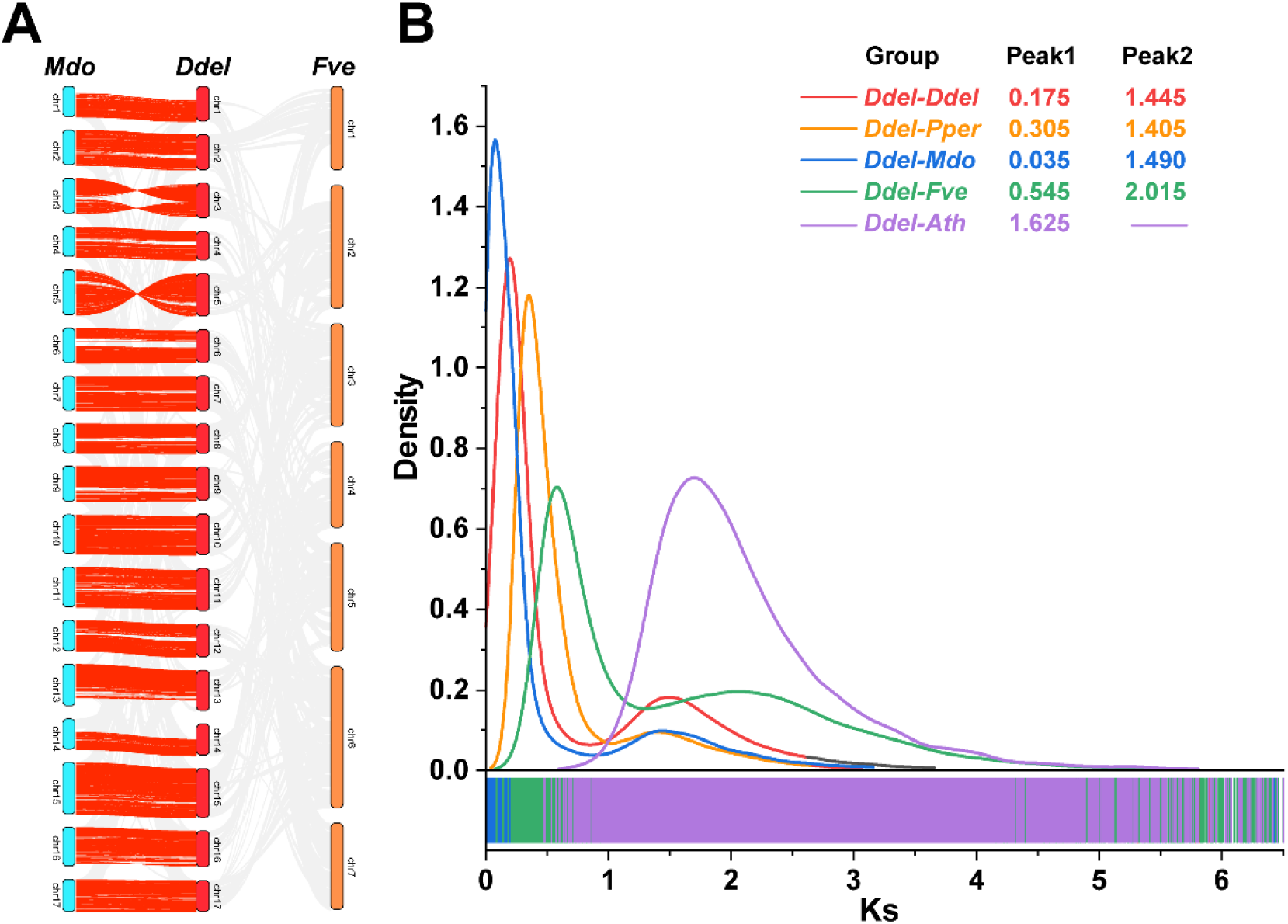
Evidence for WGD events in *D. delavayi*. A. Synteny blocks of *Mdo-Ddel-Fve*. B. Ks density distributions. *Ddel-Ddel* orthologues (red line), *Ddel-Pper* orthologues (orange line), *Ddel-Mdo* orthologues (Blue line), *Ddel-Fve* orthologues (green line) and *Ddel-Vvin* orthologues (purple line). The horizontal coordinate is the Ks value of the orthologue gene pairs, and the vertical coordinate is the density of Ks value. The abbreviations of the *Atha, Ddel, Fve, Mdo* and *Pper* represented *Arabidopsis thaliana, Docynia delavayi, Fragaria vesca, Malus domestica* and *Prunus persica*, respectively.

### Volatile organic compounds in D. delavayi fruit

The OPLS-DA demonstrated that the first two principal components explained 19.2% (PC1) and 17.4% (PC2), indicating that the discrepancy of VOCs content at different storage periods (Fig. 4A). A total of 120 VOCs were detected in *D. delavayi* fruit by HS-SPME-GC/MS, which were mainly divided into 8 categories, including esters, alkanes, terpenes, alcohols, ketones, aldehydes, ethers and other compounds. Among them, esters (36.7%) were the most abundant types VOCs in *D. delavayi* fruit, followed by aldehydes (14.2%) (Fig. 4B). Generally, the relative content of total VOCs showed an upward accumulation trend, from 4.7091μg/g at S1 to 13.1874μg/g at S4 (Fig. 4C). The relative content of different types of VOCs displayed dynamic change, of which the terpenes are the main VOCs in all storage periods, followed by esters and alkanes (Fig. 4D). Additionally, it could clearly show that the total terpene content was increased with the increasing of storage time. Further analysis for the content of five terpenes revealed that α-farnesene was the highest terpenes content, reaching a maximum of 10.53μg/g at S4, accounting for 79.9% of total VOCs content (Table S16; Fig. 4E). Hence, we considered α-farnesene to be the major VOC emitted during the *D. delavayi* fruit storage and were mining genes related to α-farnesene biosynthesis in our next study.

**Fig. 4.**
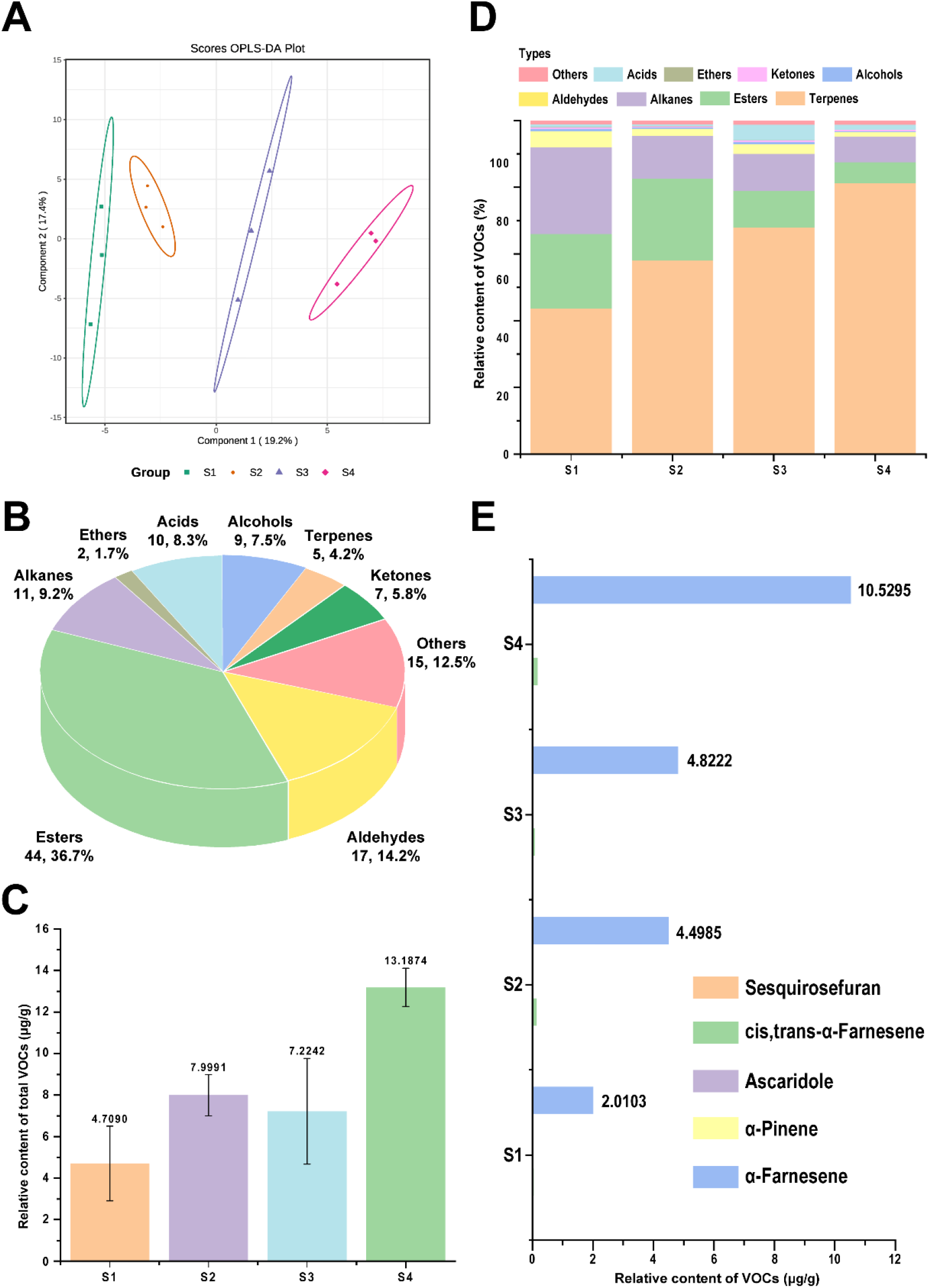
The volatile metabolome analysis of *D. delavayi* fruit at different storage periods. A. The partial least squares discriminant analysis (PLS-DA) of 12 samples at 4 storage periods. B. The statistic of VOCs types detected in *D. delavayi* fruit at 4 storage periods. C. The relative content of total VOCs at 4 storage periods. D. The percentage (%) of each type of VOCs in *D. delavayi* fruit at 4 storage periods. E. The relative content of five terpenes at 4 storage periods. S1, S2, S3 and S4 represented 0, 3, 6, 9 days after harvest, respectively. The data was mean ± SD of three biological replicates of different samples.

### Transcriptome sequencing and DEGs analysis

After filtering out data containing adapter and low-quality reads, about 72.6Gb transcriptomic data were obtained, with an average of 6.1Gb per sample. The Q20 and Q30 each sample were all above 97.92% and 94.13%, respectively, and the GC content ranged from 45.81% to 47.03%. The result of the alignment with *D. delavayi* genome showed that more than 94.59% of clean reads in each sample were successfully mapped to the reference genome, of which over 85.16% of them in each sample can be mapped to genes. Generally, the transcriptomic data was high quality and reliable in this study (Table S17). The PCA result exhibited that the first two principal components explained 55% (PC1) and 32% (PC2), indicating that the discrepancy of gene expression at different storage periods (Fig. 5A). The DEGs analysis displayed that a total of 5880 DEGs were identified in three comparison groups (S1_Vs_S2, S1_Vs_S3, and S1_Vs_S4), of which 3234 and 2646 genes were down-regulated and up-regulated expression, respectively. 570 DEGs (314 down-regulated and 256 up-regulated) were identified in S1_Vs_S2 comparison group; 3495 DEGs (1740 down-regulated and 1755 up-regulated) were identified in S1_Vs_S3 comparison group; 3992 DEGs (2492 down-regulated and 1500 up-regulated) were identified in S1_Vs_S4 comparison group. Additionally, a total of 163 DEGs were identified in all three comparison groups (Fig. 5B, C). Meanwhile, a total of 370 differential expression TFs were identified, of which ERF (36) TF family had largest number of members, followed by bHLH (31), C2H2 (29), MYB (27) and WRKY (25) TF family, and 5 TFs showed differential expression in all three groups (Table S18; Fig. 5D).

**Fig. 5.**
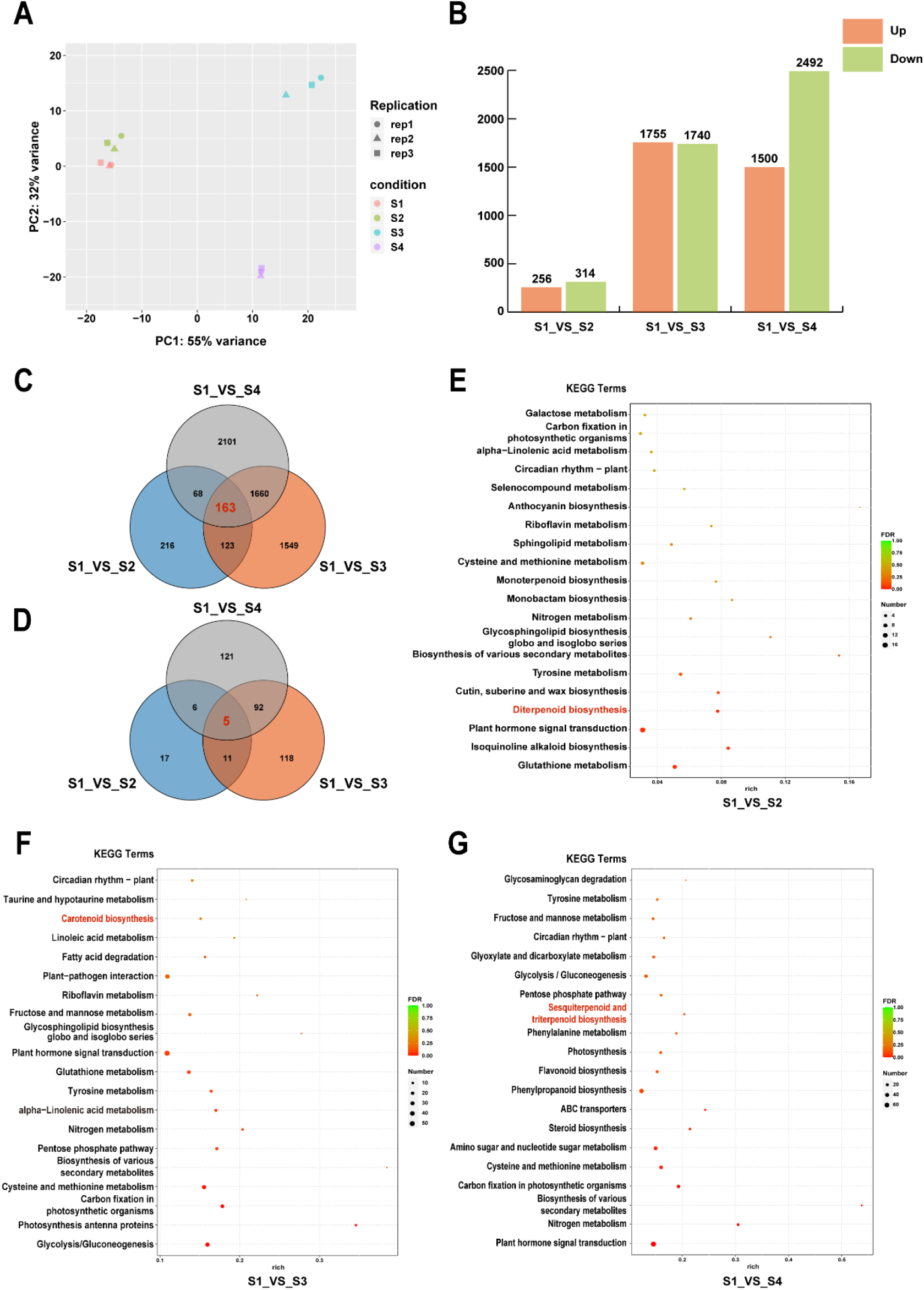
The transcriptome analysis of *D. delavayi* fruit at different storage periods. A. The principal component analysis (PCA) of 12 samples at 4 storage periods. B. The number of up-regulated and down-regulated DEGs in S1_vs_S2, S1_vs_S3 and S1_vs_S4 groups. C. Venn diagram of DEGs for S1_vs_S2, S1_vs_S3 and S1_vs_S4 groups. D. Venn diagram of TFs for S1_vs_S2, S1_vs_S3 and S1_vs_S4 groups. E∼F. The top 20 enrichment terms annotated by KEGG database of DEGs for S1_vs_S2, S1_vs_S3 and S1_vs_S4 groups. The red terms were the pathways related to ester and terpene biosynthesis. S1, S2, S3 and S4 represented 0, 3, 6, 9 days after harvest, respectively.

The DEGs were standardized and clustered to analyze their expression patterns using K-means, and the result showed that they were categorized 7 subclasses based on their expression levels at different storage periods (Fig. S7). Among them, subclass 4 (1336) was enriched with the most DEGs, followed by subclass 2 (1332) and 7 (1243). The subclass 3, 4 and 5 displayed decreasing expression patterns, while subclass 7 presented increasing expression pattern. A total of 1109 significant enriched GO terms (*P<0*.*05*) were annotated, of which response to jasmonic acid (GO:0009753), response to stimulus (GO:0050896) and response to abiotic stimulus (GO:0009628) were the most significantly enriched terms in S1_Vs_S2, S1_Vs_S3 and S1_Vs_S4 comparison groups, respectively (Fig S8). The KEGG annotation result of DEGs showed that Diterpene biosynthesis (ko00904) and Sesquiterpene and triterpene biosynthesis (ko00909) pathways were significant enriched, which related to esters and terpene biosynthesis (Fig. 5E∼G).

### Identification of α-farnesene biosynthesis pathway genes

A sesquiterpene, α-farnesene, biosynthesized through the MVA pathway, was the first aromatic components formed in *D. delavayi* fruit after harvest (Fig. 4D). Here, a hydroxymethylglutaryl-CoA reductase (HMGCR), a mevalonate kinase (MVK) and an *AFS* genes in MVA pathway were identified as differential structural genes (Fig 6A). The result of correlation analysis between α-farnesene and structural genes in MVA pathway showed that only *DdeAFS* gene exhibited a significant positive correlation (R=0.87, *P*=0.000254) with α-farnesene. Additionally, a total of 15 TFs exhibited a significant positive correlation (R>0.8, *P*<0.01) with *DdeAFS* gene and a total of 12 TFs presented a significant positive correlation (R>0.8, *P*<0.01) with the content of α-farnesene, including C2H2 (3), NAC, MYB (2), GRAS (2), bHLH (1), ERF (1), TCP (1), HD-ZIP (1), C3H (1) and NF-YA (1) TFs (Fig. 6C).

**Fig. 6.**
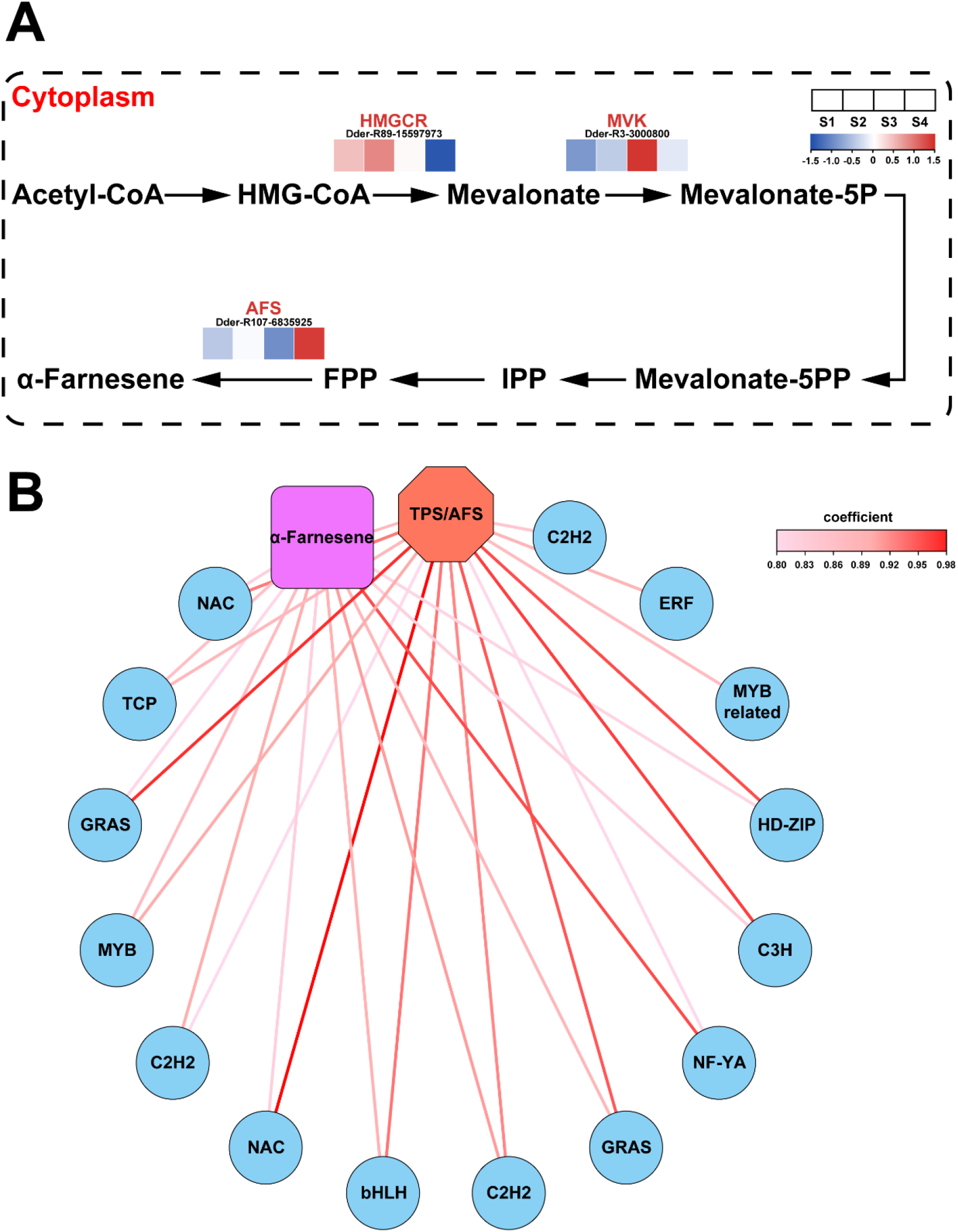
Identification of genes related to α-farnesene through KEGG pathway annotation and correlation analysis. A. MVA pathway. Red letters were the DEGs identified in this study, and the heatmaps showed their expression levels at 4 storage periods. From blue to red in the heatmap showed the expression levels of genes varying from low to high. B. Network diagram of correlation between the relative content of α-farnesene, the expression levels of DdeAFS and TFs. S1, S2, S3 and S4 represented 0, 3, 6, 9 days after harvest, respectively.

The RT-qPCR result displayed that the expression patterns of these genes were consistent with the result of RNA-seq data generally (Fig. 7A). To further investigate the function of TFs and *DdeAFS* related to α-farnesene biosynthesis, the subcellular localization and dual-luciferase assay were performed. The result showed that *DdeAFS* was ubiquitously positioned in cell membrane, cytoplasm and nucleus, and DdebHLH, DdeERF1, DdeC2H2, DdeMYB were localized in nucleus and acted as transcription activator (Fig. 7B). The dual-luciferase assay displayed that DdebHLH, DdeERF1, DdeMYB were able to significantly improve the activity of the *DdeAFS* promotor by compared with the control group, suggesting that they were positive regulators of *DdeAFS* expression. However, there was no significant discrepancy between DdeC2H2 and control group, indicating that *DdeAFS* was not the downstream target gene of DdeC2H2 (Fig. 7D).

**Fig. 7.**
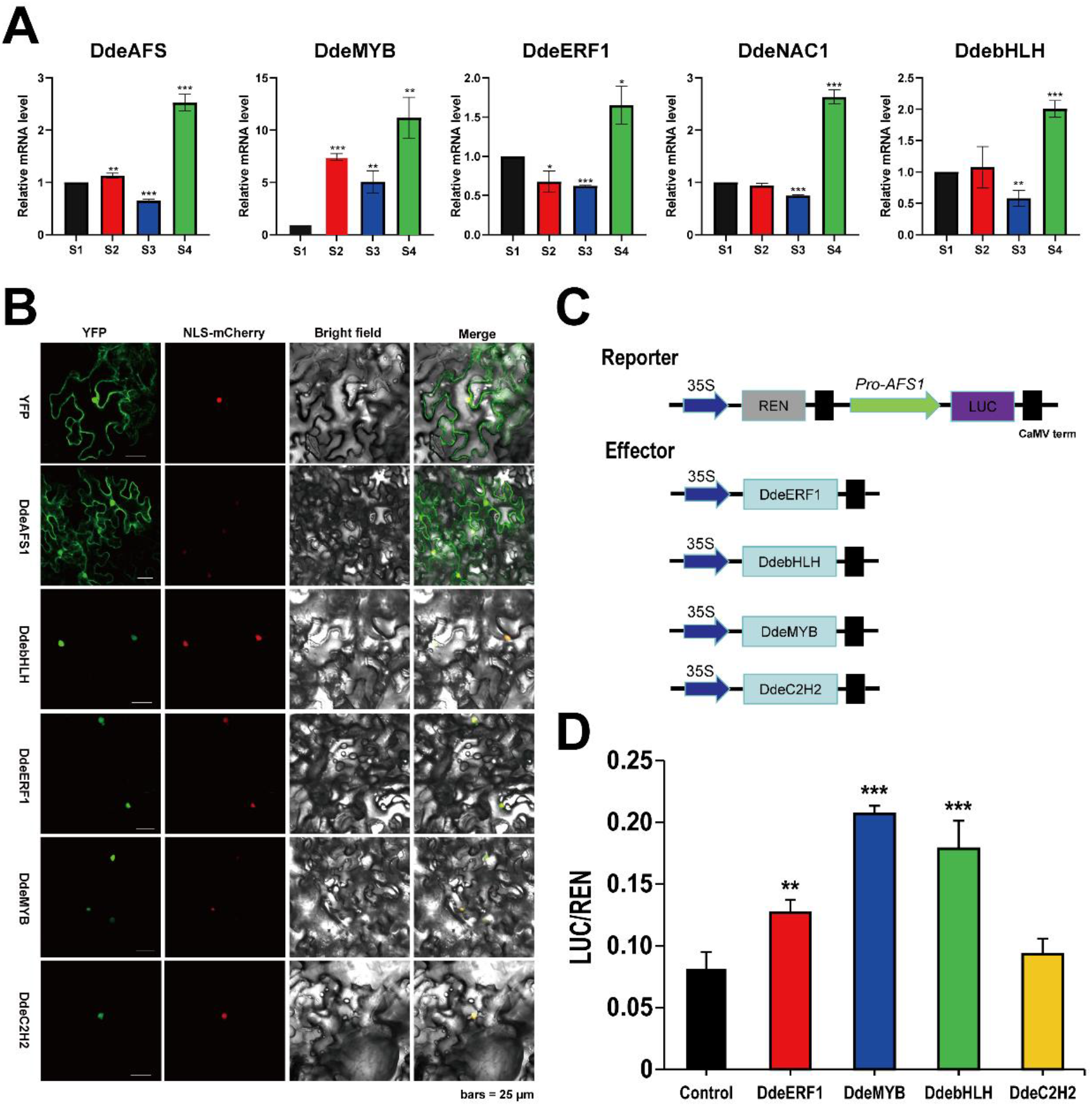
The functional assay of candidate gene in α-farnesene biosynthesis. A. The relative expression level of 5 random candidate genes through the RT-qPCR. The relative expression level of genes in S1 was normalized as 1 and regarded as the control group to perform the significance analysis. B. Subcellular localization analysis of candidate genes in α-farnesene biosynthesis. C. The structural diagram of reporter and effector plasmids. D. The transcriptional activity of DdeERF1, DdeMYB, DdebHLH and DdeC2H2 effectors. Statistical significance was determined by One-way ANOVA and Tukey’s tests. Asterisks indicated a significant difference between the treatment and control groups. S1, S2, S3 and S4 represented 0, 3, 6, 9 days after harvest, respectively. Error bars represent the mean ± SD of three biological replicates.

## Discussion

Fruit trees are high heterogeneity in generally, making them more challenging to assemble high-quality complete genome (Jiang et al., 2019). Consistent with most fruit trees, the high heterozygosity (0.54%) and repeat rate (23.7%) in *D. delavayi* genome presented challenges to genome assembly (Huang et al., 2021; Jiang et al., 2019; Xia et al., 2021). Compared with other sequenced species of the Rosaceae family, both the scaffold N50 (35.15 Mb) and anchoring rate (93.8%) of *D. delavayi* genome were at higher levels, indicating that the quality of *D. delavayi* genome assembly was better (Chen et al., 2021a; Jiang et al., 2019; Jiu et al., 2023). 96.5% complete genes were identified through the BUSCO assessment analysis, which further demonstrated that the *D. delavayi* genome was a high-quality and was available for the subsequent analysis. The TE content is one of the major factors of plant genome size, and it is positively correlated with genome size (Wendel et al., 2016). The conclusion remains tenable by comparing the genome data of all sequenced Rosaceae family plant (Jung et al., 2019).

WGD events in the evolutionary history of angiosperms have greatly contributed to plant adaptation in the changing global environment, with almost all angiosperms undergoing at least a WGD event (Meng et al., 2023; Qin et al., 2023). In this study, a total of two WGD events were identified in *D. delavayi* genome, corresponding to a recent (Ks=0.175) and a Paleo-WGD (Ks=1.445) event, respectively. Since the recent WGD event in *D. delavayi* occurred before it diverged from *M. domestica*, we inferred that the recent WGD event might have been shared with *M. domestica* (Velasco et al., 2010). Meanwhile, the collinearity of *D. delavayi* large inter-chromosomal segments and its well-collinearity with *M. domestica* could also support this genome duplication pattern (Daccord et al., 2017). Taken together, it inferred that the formation of 17 chromosomes in *D. delavayi* might also conform to the autopolyploidization hypothesis as same as *M. domestica* (Evans and Campbell, 2002; Velasco et al., 2010). The chromosomes of *D. delavayi* and *M. domestica* displayed a well-collinearity relationship, suggesting that the numbers of chromosome were conserved with few rearrangements after the divergence of *D. delavayi* and *M. domestica* (Zhang et al., 2021). Furthermore, compared the Ks value for *D. delavayi* Paleo-WGD event with that in the Rosaceae plants, we found that the Paleo-WGD event might be shared by the Rosaceae family (Wang et al., 2021).

The evolutionary relationship of *D. delavayi* was resolved using its genome information, and the result equally supported the conclusion that *M. domestica* and *D. delavayi* were more closely related (Li et al., 2022b). The divergence time between *M. domestica* and *D. delavayi* was approximately 11MYA, which was consistent with the result of the Ks analysis, indicating that the generation of *D. delavayi* originated form the divergence of *M. domestica*. Additionally, combined with previous research we found that the *M. domestica* might be the common ancestor of all species with 17 chromosomes in Rosaceae family (Chen et al., 2019; Jiang et al., 2020). The contraction and expansion gene families are key drives of adapting natural variation and phenotypic diversity in plants (Chen et al., 2021b; Chen et al., 2013). According to the phylogenetic result of 10 species, 2254 and 1557 gene families were identified as contraction and expansion gene families in *D. delavayi* genome, respectively. The phenomenon of fewer expanded and more contracted gene families was also commonly found in other Rosaceae plants, suggesting that they have undergone similar evolutionary patterns (Qin et al., 2023). Notably, as with most aromatic plants, the expansion of terpene-related gene families in *D. delavayi* genome was one of the key factors that gave them special aroma (Dong et al., 2021; Huang et al., 2021; Shen et al., 2023).

Aroma is a critical trait of fruit flavor and quality, which usually undergoes dynamic changes during fruit post-harvest ripening period due to a series of biochemical reactions, especially in respiratory climacteric fruits (Liu et al., 2021a; Wang et al., 2022a). It has reported that aroma discrepancy existed between *D. delavayi* fruits of different maturity levels (Wang et al., 2023b), while our resource investigations revealed that it became more significant during post-harvest storage period. In this study, the total aroma content displayed a gradual upward trend during fruit post-harvest storage period, of which α-farnesene was the most abundant VOCs. α-Farnesene was released mainly after fruit ripening, which might contribute to plant defense and attract animals for spreading *D. delavayi* seeds (Pechous and Whitaker, 2004; Wang et al., 2020). In addition, temperature was also one of the crucial factors for the α-farnesene content in fruit, with low-temperatures during fruit storage accelerating its accumulation (Pechous and Whitaker, 2004). In summary, the contents of aroma components during *D. delavayi* fruit post-harvest period underwent significant changes, of which α-farnesene was the dominant VOCs, imparting a strong and specific aroma to *D. delavayi* fruit.

α-Farnesene biosynthesis was mainly affected by the *AFS* genes in plant, and its expression was generally synchronous with its product (Du et al., 2022; Xiang et al., 2022). In this study, a *DdeAFS* gene was identified, which showed a significantly positive correlation with α-farnesene content, inferring that it was a key enzyme gene in α-farnesene biosynthesis in *D. delavayi* fruit. Consistent with previous studies, *DdeAFS* was localized in the cytoplasm and involved in α-farnesene biosynthesis via the cytoplasmic-located MVA pathway (Vranová et al., 2013; Zhang et al., 2019). Differently, it was also localized in the cell nucleus and membrane, speculating that the result might be related to its functional diversity (Zhang et al., 2022). Pervious studies have reported that both positive and negative regulators of *AFS* gene expression were presented in apple (Du et al., 2022; Wang et al., 2020), and these types of TFs were also preliminarily screened as potential regulators for α-farnesene biosynthesis in this study. Among them, overexpression of MdERF3 and MdMYC2 in apple fruit significantly up-regulated the expression of *MdAFS*, thereby increased the production of α-farnesene (Wang et al., 2020). Similarly, DdeERF and DdebHLH were also identified as the candidate gene in regulating α-farnesene biosynthesis, which could significantly activate the promoter activity of *DdeAFS* gene, further demonstrating their potential role in α-farnesene biosynthesis in *D. delavayi* fruit. MdLSD1, a TF encoded C2H2 type proteins, played a negative role in regulating α-farnesene biosynthesis at transcription level in apple fruit (Du et al., 2022). Differently, the DdeC2H2 TF initially screened in this study failed to significantly change the promoter activity of *AFS* gene, which was presumed to be caused by structural differences between DdeC2H2 and MdLSD1 (Du et al., 2022; Jiao et al., 2020).

In addition, some potential TFs related to α-farnesene biosynthesis were also identified in the present study, such as MYB, NAC, HD-ZIP etc. Among them, MYB is one of vital TF families in plant genome, which is widely involved in plant various life processes, especially in regulating the secondary metabolic biosynthesis (Cao et al., 2020). Pervious studies have mainly concentrated on their involvement in the regulation of flavonoids, anthocyanin and lignins biosynthesis (Cao et al., 2020). However, with the emphasis on fruit flavor substances, increasingly studies have begun to focus on their role in regulating fruit aroma substances biosynthesis in recent years (Li et al., 2023; Medina-Puche et al., 2015, Zhang et al., 2018). The roles of MYB TFs in regulating certain sesquiterpenes biosynthesis have been identified in some plants (Abbas et al., 2021, Bedon et al., 2010), while their role in modulating α-farnesene biosynthesis has not been reported to our knowledge. In this study, a MYB TF (DdeMYB) involved in the positive regulation of α-farnesene biosynthesis in *D. delavayi* fruit was first identified. It showed a significantly positive correlation with *DdeAFS* expression and α-farnesene content, and it could also significantly activate the promoter activity of the *DdeAFS*. These results suggested that DdeMYB could regulated α-farnesene biosynthesis through activating the *DdeAFS* expression in *D. delavayi* fruit, which provided a new insight into the functional analysis of MYB TFs.

### Conclusion

In this study, a 612.98 Mb chromosome-scale genome was first assembled using the combination of ONT sequencing and Hi-C scaffolding technology. Then, a *DdeAFS* gene and 15 TFs potentially involved in α-farnesene biosynthesis were selected through the combination of genomic, transcriptomic and metabolomic analysis. Lastly, three TFs were identified as the key regulator in α-farnesene biosynthesis, of which it was noteworthy that our study found a novel function for MYB TF in regulating the transcription of the *AFS* gene through the dual-luciferase assay. Thus, this work not only provided high-quality genomic information for *D. delavayi*, but also contributed to its genetic improvement of important economic traits.

## Materials and Methods

### Genome sequencing, assembly and quality assessment

The healthy leaf samples of *D. delavayi* were collected from Lancang, Yunnan province of China to perform genomic DNA (gDNA) isolation. The gDNA purity was detected using Nanodrop, and then the Qubit was then used to accurately quantify the gDNA. After the gDNA was qualified, the whole genome sequencing was commissioned by the BGI, Shenzhen, China. The jellyfish program was used for the K-mer analysis (K=17) to estimate the genome size and heterozygosity, and the GenomeScope was used to plot the K-mer distribution (Marçais and Kingsford, 2011, Vurture et al., 2017).

After determining the genome size, the first assembly of the genome was performed as described by Xia et al (2021). Three *de novo* error correction of the assembly result was first conducted using the Racon, then the assembly result was compared with the second-generation sequencing data to correct errors with 5 rounds, thus eventually obtaining the assembly result (Senol-Cali et al., 2019). Then, Benchmarking Universal Single-Copy Orthologs (BUSCO) was used to assess the genomic quality. Furthermore, to obtain a chromosome-scale assembly, gDNA was extracted for constructing the Hi-C library, and the detailed steps referred to Wang et al (2021).

### Genome annotation

The repetitive sequences contain both types of tandem repeat (TR) and transposable elements (TEs). The TR elements were searched using the TRF software (Benson, 1999). The TEs were annotated using homologous prediction and *de novo* prediction, and the method was consistent with the description of Liu et al (2023). The structural annotation of *D. delavayi* were carried out through de novo, homologous and transcript-based prediction (Shen et al., 2023). The information of exons, introns, CDS and protein-coding genes was compared with those of closely related species (*M. domestica, Pyrus communis, Rosa chinensis* and *Fragaria vesca*) and the distribution chart was drawn. The tRNA was identified using the tRNAscan-SE software, and the miRNA, rRNA and snRNA were annotated using Rfam database. The gene function was annotated using NR, GO, KEGG, KOG, TrEMBL, Interpro and Swissprot with default parameters.

### Analysis of gene families and phylogenetic evolution

Five representative species in Rosaceae family (*M. domestica, P. communis, Prunus persica, Rosa rugosa* and *F. vesca*), three representative fruit species (*Solanum lycopersicum, Vitis vinifera* and *Ziziphus jujuba*), two model plants (*Populus trichocarpa* and *Arabidopsis thaliana*) and a monocotyledon plant (*Oryza sativa*) were selected for comparative genomic analysis (Table S1). The ortholog clusters of protein-coding genes in *D. delavayi* and 11 other species were identified referred to previous study (Liu et al., 2023). Referring to the method of pervious study (Liu et al., 2023), we selected the single-copy genes of *D. delavayi* and the nine species mentioned above to construct the phylogenetic tree, identify the expansion and contraction gene families, and predict the divergence time. Last, the GO and KEGG databases was used to perform the functional annotation of the expansion gene families.

### Analysis of genome synteny and WGD

To analyze the syntenic relationship between *D. delavayi*, and *A. thaliana, M. domestica, F. vesca* and *P. persica*, the MCScanX program was used to found the genome synteny blocks with a minimum of 30 genes in syntenic region and the TBtools software was used to draw the chromosomal synteny map (Chen et al., 2020, Wang et al., 2012). The Ks value of collinear gene pairs in *D. delavayi*-*D. delavayi, D. delavayi*-*M. domestica, D. delavayi*-*P. persica, D. delavayi*-*F. vesca* and *D. delavayi*-*A. thaliana* was analyzed via the Ka/Ks calculator program of TBtools software and the distribution chart of Ks density were drawn (Chen et al., 2020).

### Samples preparation and VOCs detection

A total of 60 ripe fruits (180 days after flowering) with similar size and without visible defects were harvested from Lancang county at maturity (180 days after flowering), and were sent to the laboratory for storage under natural conditions. Samples were taken every three days, with a total of four sampling periods (0, 3, 6 and 9 days after harvest, denoting by S1, S2, S3 and S4 in subsequent analyses, respectively). Every five fruits were mixed into one biological replicate, with a total of 3 biological replicates per sampling period. Samples were rinsed with sterile water and frozen in liquid nitrogen immediately for VOCs detection and RNA extraction. The conditions of HS-SPME and GC-MS were consistent with the description of Wang et al (2023b).

### RNA extraction, sequencing and transcriptome analysis

The extraction of total RNA and transcriptome sequencing were entrusted to Shanghai Bioprofile Technology Company Ltd. (Shanghai, China). The Illumina HiSeq 6000 platform was used to transcriptome sequencing, and the high-quality reads were mapped to the *D. delavayi* genome data. The gene function was annotated via the GO and KEGG database. The quantification of transcript abundance and identification of differential expression genes (DEGs) were referred to the previous researches (Wang et al., 2022b; Xia et al., 2021).

### Identification of terpenes biosynthesis pathway genes

The correlation network analysis between differential structural genes, TFs and VOCs was conducted using the Metware Cloud., The differential structural genes, TFs and VOCs with correlation coefficient R>0.8 and *P*<0.05 were chosen to draw correlation networks using Cytoscape software (Doncheva et al., 2019).

### RT-qPCR, subcellular localization and dual-luciferase assay

Five genes were randomly selected for RT-qPCR assay to verify the expression of genes in this study. Both the housekeeping gene and reaction system referred to the study of Ding et al (2021). The Primer 5.0 software was used to design the primer sequences of the five genes above (Table S2). To determine the subcellular localizations of *DdeAFS*, DdebHLH, DdeERF1, DdeMYB and DdeC2H2 proteins, coding regions of relevant genes were fused in-frame to the upstream of the YFP sequence in the pCambia1300-35S-YFP vector, respectively. The resulting constructs were co-transfected with nuclear marker pCambia-35S-NLS-mCherry into the tobacco (*Nicotiana tabacum*) leaves. After being incubated in dark for two days, fluorescence of YFP and mCherry in the leaf epidermal cells was imagined by a confocal laser scanning microscope (TCS SP8, Leica), excitation and emission wavelengths of 520 and 550 nm for YFP, 575 and 650 nm for mCherry.

pGreenII-0800-LUC recombinant reporter plasmid (with a 2-kb promoter of *AFS* gene) and pGreenII 62-SK recombinant effector plasmids (with transcriptional factor DdeERF1, DdeMYB, DdebHLH or DdeC2H2) were transformed into *Agrobacterium tumefaciens* GV3101, with the pGreenII constructs being co-transformed with a pSoupP19 plasmid. Combinations of reporter and effectors were co-infiltrated into leaves of *N. benthamiana* and the luciferase activity of the tobacco extracts was analyzed using a Promega Dual-Luciferase Assay Kit and detected on a Bio-Tek Synergy 2 multimode microplate reader.

### Statistical analysis

The principal component analysis (PCA) and orthogonal partial least squares discriminant analysis (OPLS-DA) were performed using the R package. SPSS v21 software was used to perform the one-way ANOVA tests at least three biological replicates (*P*<0.05). Results were exhibited as the mean ± standard deviation.

## Acknowledgments

This work is supported by the National Natural Science Foundation of China (No. 32060350), Key Basic Research Projects in Yunnan Province (202401AS070042), Forestry Innovation Program of Southwest Forestry University (Grant: LXXK-2023Z01), and the Fund of Ten-Thousand Talent Plans for Young Top-notch Talents of Yunnan Province [Grant No. YNWR–QNBJ–2020-230].

## Author contributions

**Jinhong Tian**: Conceptualization; Data curation; Formal analysis; Investigation; Methodology; Writing-review and editing. **Zhuo Chen**: Conceptualization; Data curation; Formal analysis; Investigation; Methodology; Writing-review and editing. **Can Jiang**: Formal analysis; Investigation; **Siguang Li**: Investigation; Methodology. **Xinhua Yun**: Conceptualization; Resources; Data curation; Supervision; Writing-original draft; Writing-review and editing. **Chengzhong He**: Conceptualization; Resources; Data curation; Supervision; Writing-original draft; Writing-review and editing. **Dawei Wang**: Conceptualization; Resources; Data curation; Formal analysis; Supervision; Funding acquisition; Writing-original draft; Project administration; Writing-review and editing.

## References

Abbas F, Ke YG, Zhou YW, Yu YY, Waseem M, Ashraf U, Wang CT, Wang XY, Li XY, Yue YC, Yu RC, Fan YP (2021) Genome-wide analysis reveals the potential role of MYB transcription factors in floral scent formation in Hedychium coronarium. Front Plant Sci 12: 623742

Bedon F, Bomal C, Caron S, Levasseur C, Boyle B, Mansfield SD, Schmidt A, Gershenzon J, Grima-Pettenati J, Séguin A, MacKay J (2010) Subgroup 4 R2R3-MYBs in conifer trees: gene family expansion and contribution to the isoprenoid- and flavonoid-oriented responses. J Exp Bot 61: 3847–3864

Benson G (1999) Tandem repeats finder: a program to analyze DNA sequences. Nucleic Acids Res 27: 573–580

Cao YP, Li K, Li YL, Zhao XP, Wang LH (2020) MYB transcription factors as regulators of secondary metabolism in plants. Biology 9: 61

Chen C, Xia X, Wang DW (2022) Identification of nutritional components in unripe and ripe Docynia delavayi (Franch.) Schneid fruit by widely targeted metabolomics. PeerJ 10: e14441

Chen CJ, Chen H, Zhang Y, Thomas HR, Frank MH, He YH, Xia R (2020) TBtools: an integrative toolkit developed for interactive analyses of big biological data. Mol Plant 13: 1194–1202

Chen F, Su LY, Hu SY, Xue JY, Liu H, Liu GH, Jiang YF D. JK, Qiao YS, Fan YN, Liu H, Yang Q, Lu WJ, Shao ZQ, Zhang J, Zhang LS, Chen F, Cheng ZMM (2021a) A chromosome-level genome assembly of rugged rose (Rosa rugosa) provides insights into its evolution, ecology, and floral characteristics. Hortic Res 8: 141

Chen JY, Xie FF, Cui YZ, Chen CB, Lu WJ, Hu XD, Hua QZ, Zhao J, Wu ZJ, Gao D, Zhang ZK, Jiang WK, Sun QM, Hu GB, Qin YH (2021b) A chromosome-scale genome sequence of pitaya (Hylocereus undatus) provides novel insights into the genome evolution and regulation of betalain biosynthesis. Hortic Res 8: 164

Chen SD, Krinsky BH, Long MY (2013) New genes as drivers of phenotypic evolution. Nat Rev Genet 14: 646–660

Chen X, Li S, Zhang D, Han M, Jin X, Zhao C, Wang S, Xing L, Ma J, Ji J, An N (2019) Sequencing of a wild apple (Malus baccata) genome unravels the differences between cultivated and wild apple species regarding disease resistance and cold tolerance. G3-Genes Genom Genet 9: 2051–2060

Daccord N, Celton JM, Linsmith G, Becker C, Choisne N, Schijlen E, van de Geest H, Bianco L, Micheletti D, Velasco R, Di Pierro EA, Gouzy J, Rees DJG, Guérif P, Muranty H, Durel CE, Laurens F, Lespinasse Y, Gaillard S, Aubourg S et al. (2017) High-quality de novo assembly of the apple genome and methylome dynamics of early fruit development. Nat Genet 49: 1099–1106

Deng XK, Zhao XP, Lan Z, Jiang J, Yin W, Chen LY (2014) Anti-tumor effects of flavonoids from the ethnic medicine Docynia delavayi (Franch.) Schneid. and its possible mechanism. J Med Food 17: 787–794

Ding RR, Che XK, Shen Z, Zhang YH (2021) Metabolome and transcriptome profiling provide insights into green apple peel reveals light- and UV-B-responsive pathway in anthocyanins accumulation. BMC Plant Biol 21: 351

Doncheva NT, Morris JH, Gorodkin J, Jensen LJ (2019) Cytoscape stringapp: network analysis and visualization of proteomics data. J Proteome Res 18: 623–632

Dong SS, Liu M, Liu Y, Chen F, Yang T, Chen L, Zhang XT, Guo X, Fang DM, Li LZ, Deng T, Yao ZX, Lang XA, Gong YQ, Wu E, Wang YL, Shen YM, Gong X, Liu H, Zhang SZ (2021) The genome of Magnolia biondii Pamp. provides insights into the evolution of Magnoliales and biosynthesis of terpenoids. Hortic Res 8: 38

Du B, Ma X, Liu H, Dong K, Liu H, Zhang Y (2022) Transcription factor MdLSD1 negatively regulates α-farnesene biosynthesis in apple-fruit skin tissue. Plant Biol 24: 1076–1083

Evans RC, Campbell CS (2002) The origin of the apple subfamily (Maloideae; Rosaceae) is clarified by DNA sequence data from duplicated GBSSI genes. Am J Bot 89: 1478–1484

Huang C, Sun PX, Yu S, Fu GY, Deng Q, Wang ZW, Cheng SH (2023) Analysis of volatile aroma components and regulatory genes in different kinds and development stages of pepper fruits based on non-targeted metabolome combined with transcriptome. Int J Mol Sci 24: 7901

Huang ZY, Shen F, Chen Y, Cao K, Wang LR (2021) Chromosome-scale genome assembly and population genomics provide insights into the adaptation, domestication, and flavonoid metabolism of Chinese plum. Plant J 108: 1174–1192

Initiative IPG, Verde I, Abbott AG, Scalabrin SQ, Jung S, Shu S, Marroni F, Zhebentyayeva T, Dettori MT, Grimwood J, Cattonaro F, Zuccolo A, Rossini L, Jenkins J, Vendramin E, Meisel LA, Decroocq V, Sosinski B, Prochnik S, Mitros T et al. (2013) The high-quality draft genome of peach (Prunus persica) identifies unique patterns of genetic diversity, domestication and genome evolution. Nat Genet 45: 487–494

Jiang FC, Zhang JH, Wang S, Yang L, Luo YF, Gao SH, Zhang ML, Wu SY, Hu SN, Sun HY, Wang YZ (2019) The apricot (Prunus armeniaca L.) genome elucidates Rosaceae evolution and beta-carotenoid synthesis. Hortic Res 6: 128

Jiang S, An HS, Xu FJ, Zhang XY (2020) Chromosome-level genome assembly and annotation of the loquat (Eriobotrya japonica) genome. Gigascience 9: giaa015

Jiao ZC, Wang LP, D. H, Wang Y, Wang WX, Liu JJ, Huang JH, Huang W, Ge LF (2020) Genome-wide study of C2H2 zinc finger gene family in Medicago truncatula. BMC Plant Biol 20: 401

Jiu S, Chen B, Dong X, Lv Z, Wang Y, Yin C, Xu Y, Zhang S, Zhu J, Wang J, Liu X, Sun W, Yang G, Li M, Li S, Zhang Z, Liu R, Wang L, Manzoor MA, José QG et al. (2023) Chromosome-scale genome assembly of Prunus pusilliflora provides novel insights into genome evolution, disease resistance, and dormancy release in Cerasus L. Hortic Res 10: uhad062

Jung S, Lee T, Cheng CH, Buble K, Zheng P, Yu J, Humann J, Ficklin SP, Gasic K, Scott K, Frank M, Ru S, Hough H, Evans K, Peace C, Olmstead M, DeVetter LW, McFerson J, Coe M, Wegrzyn JL et al. (2019) 15 years of GDR: New data and functionality in the Genome Database for Rosaceae. Nucleic Acids Res 47: D1137–D1145

Li LX, Fang Y, Li D, Zhu ZH, Zhang Y, Tang ZY, Li T, Chen XS, Feng SQ (2023) Transcription factors MdMYC2 and MdMYB85 interact with ester aroma synthesis gene MdAAT1 in apple. Plant Physiol 193: 2442–2458

Li LX, Ou WL, Wang YC, Peng JY, Wang DW, Xu S (2022a) Comparison of genetic diversity between ancient and common populations of Docynia delavayi (Franch.) Schneid. gene 829: 146498

Li LX, Peng JY, Wang DW, Duan AA (2022b) Chloroplast genome phylogeny and codon preference of Docynia longiunguis. Chinese Journal of Biotechnology 38: 328–342

Liu MM, Li C, Jiang T, Wang RP, Wang Y, Zhang WE, Pan XJ (2023) Chromosome-scale genome assembly provides insights into flower coloration mechanisms of Canna indica. Int J Biol Macromol 251: 126148

Liu XJ, Hao NN, Feng RF, Meng ZP, Li YN, Zhao ZY (2021a) Transcriptome and metabolite profiling analyses provide insight into volatile compounds of the apple cultivar ‘Ruixue’ and its parents during fruit development. BMC Plant Biol 21: 231

Liu YH, Wang ZX, Cui ZY, Qi QS, Hou J (2021b) α-Farnesene production from lipid by engineered Yarrowia lipolytica. Bioresources and Bioprocessing 8: 78

Marçais G, Kingsford C (2011) A fast, lock-free approach for efficient parallel counting of occurrences of k-mers. Bioinformatics 27: 764–770

Medina-Puche L, Molina-Hidalgo FJ, Boersma M, Schuurink RC, López-Vidriero I, Solano R, Franco-Zorrilla JM, Caballero JL, Blanco-Portales R, Muñoz-Blanco J (2015) An R2R3-MYB transcription factor regulates eugenol production in ripe strawberry fruit receptacles. Plant Physiol 168: 598–614

Meng FB, Chu TZ, Feng PM, Li N, Song C, Li CJ, Leng L, Song XM, Chen W (2023) Genome assembly of Polygala tenuifolia provides insights into its karyotype evolution and triterpenoid saponin biosynthesis. Hortic Res 10: uhad139

Pechous SW, Whitaker BD (2004) Cloning and functional expression of an (E, E)-alpha-farnesene synthase cDNA from peel tissue of apple fruit. Planta 219: 84–94

Peng JY, Shi C, Wang DW, Li SZ, Zhao XL, Duan AA, Cai NH, Heng CZ (2021) Genetic diversity and population structure of the medicinal plant Docynia delavayi (Franch.) Schneid revealed by transcriptome-based SSR markers. J Appl Res Med Aroma 21: 100294

Qin SJ, Xu GX, He JL, Li LJ, Ma HY, Lyu DG (2023) A chromosome-scale genome assembly of Malus domestica, a multi-stress resistant apple variety. Genomics 115: 110627

Senol-Cali D, Kim JS, Ghose S, Alkan C, Mutlu O (2019) Nanopore sequencing technology and tools for genome assembly: computational analysis of the current state, bottlenecks and future directions. Brief Bioinform 20: 1542–1559

Shen L, Ding CJ, Zhang WX, Zhang TQ, Li ZH, Zhang J, Chu YG, Su XH (2023) The Populus koreana genome provides insights into the biosynthesis of plant aroma. Ind Crop Prod 197: 116453

Velasco R, Zharkikh A, Affourtit J, Dhingra A, Cestaro A, Kalyanaraman A, Fontana P, Bhatnagar SK, Troggio M, Pruss D, Salvi S, Pindo M, Baldi P, Castelletti S, Cavaiuolo M, Coppola G, Costa F, Cova V, Dal Ri A, Goremykin V et al. (2010) The genome of the domesticated apple (Malus × domestica Borkh.). Nat Genet 42: 833–839

Vranová E, Coman D, Gruissem W (2013) Network analysis of the MVA and MEP pathways for isoprenoid synthesis. Annu Rev Plant Biol 64: 665–700

Vurture GW, Sedlazeck FJ, Nattestad M, Underwood CJ, Fang H, Gurtowski J, Schatz MC (2017) GenomeScope: fast reference-free genome profiling from short reads. Bioinformatics 33: 2202–2204

Wang AN, Peng XW, Kan H, Wang DW, Hu X, Liu Y (2023a) Extraction of flavonoids from Docynia delavayi and their antioxidant and hypoglycemic activities. Science and Technology of Food Industry 44: 232–240

Wang LJ, Lei T, Han GM, Yue JY, Zhang XR, Yang Q, Ruan HX, Gu CY, Zhang Q, Qian T, Zhang NN, Qian W, Wang Q, Pang XJ, Shu Y, Gao LP, Wang YS (2021) The chromosome-scale reference genome of Rubus chingii Hu provides insight into the biosynthetic pathway of hydrolyzable tannins. Plant J 107: 1466–1477

Wang Q, Liu H, Zhang M, Liu SH, Hao YJ, Zhang YH (2020) MdMYC2 and MdERF3 positively co-regulate α-farnesene biosynthesis in apple. Front Plant Sci 11: 512844

Wang QH, Gao F, Chen XX, Wu WJ, Wang L, Shi JL, Huang Y, Shen YY, Wu GL, Guo JX (2022a) Characterization of key aroma compounds and regulation mechanism of aroma formation in local Binzi (Malus pumila × Malus asiatica) fruit. BMC Plant Biol 22: 532

Wang Y, He YH, Liu Y, Wang DW (2023b) Analyzing volatile compounds of young and mature Docynia delavayi fruit by HS-SPME-GC-MS and rOAV. Foods 12: 59

Wang Y, Tang H, Debarry JD, Tan X, Li J, Wang X, Lee TH, Jin H, Marler B, Guo H, Kissinger JC, Paterson AH (2012) MCScanX: a toolkit for detection and evolutionary analysis of gene synteny and collinearity. Nucleic Acids Res 40: e49

Wang YC, Song YY, Wang DW (2022b) Transcriptomic and metabolomic analyses providing insights into the coloring mechanism of Docynia delavayi. Foods 11: 2899

Wendel JF, Jackson SA, Meyers BC, Wing RA (2016) Evolution of plant genome architecture. Genome Biol 17: 37

Xia X, Chen C, Yang L, Wang YC, Duan AA, Wang DW (2022) Analysis of metabolites in young and mature Docynia delavayi (Franch.) Schneid leaves using UPLC-ESI-MS/MS. PeerJ 10: e12844

Xia ZQ, Huang DM, Zhang SK, Wang WQ, Ma FN, Wu B, Xu Y, Xu BQ, Chen D, Zou ML, Xu HY, Zhou XC, Zhan RL, Song S (2021) Chromosome-scale genome assembly provides insights into the evolution and flavor synthesis of passion fruit (Passiflora edulis Sims). Hortic Res 8: 14

Xiang LE, He P, Shu GP, Yuan M, Wen M, Lan X, Liao ZH, Tang YL (2022) AabHLH112, a bHLH transcription factor, positively regulates sesquiterpenes biosynthesis in Artemisia annua. Front Plant Sci 13: 973591

Xu L, Liu Y, Yang L, Tian JH, Li JT, Ma HT, Li EL, Wang DW (2023) Plus tree selection of fresh Docynia delavayi (Franch.) Schneid. Journal of Plant Genetic Resources 24: 903–910

Zhang AD, Xiong YH, Fang J, Jiang XH, Wang TF, Liu KC, Peng HX, Zhang XJ (2022) Diversity and Functional Evolution of Terpene Synthases in Rosaceae. Plants 11: 736

Zhang QX, Chen WB, Sun LD, Zhao FY, Huang BQ, Yang WR, Tao Y, Wang J, Yuan ZQ, Fan GY, Xing Z, Han CL, Pan HT, Zhong X, Shi WF, Liang XM D. DL, Sun FM, Xu ZD, Hao RJ et al. (2012) The genome of Prunus mume. Nat Commun 3: 1318

Zhang XH, Niu MY, Teixeira da Silva JA, Zhang YY, Yuan YF, Jia YX, Xiao YY, Li Y, Fang L, Zeng SJ, Ma GH (2019) Identification and functional characterization of three new terpene synthase genes involved in chemical defense and abiotic stresses in Santalum album. BMC Plant Biol 19: 115

Zhang YX, Zhang GQ, Zhang DY, Liu XD, Xu XY, Sun WH, Yu X, Zhu XE, Wang ZW, Zhao X, Zhong WY, Chen HF, Yin WL, Huang TB, Niu SC, Liu ZJ (2021) Chromosome-scale assembly of the Dendrobium chrysotoxum genome enhances the understanding of orchid evolution. Hortic Res 8: 183

Zhang YY, Yin XR, Xiao YW, Zhang ZY, Li SJ, Liu XF, Zhang B, Yang XF, Grierson D, Jiang GH, Klee HJ, Chen KS (2018) An ETHYLENE RESPONSE FACTOR-MYB transcription complex regulates furaneol biosynthesis by activating QUINONE OXIDOREDUCTASE expression in strawberry. Plant Physiol 178: 189–201

Zhao XP, Shu GW, Chen LY, Mi X, Mei ZN, Deng XK (2012) A flavonoid component from Docynia delavayi (Franch.) Schneid represses transplanted H22 hepatoma growth and exhibits low toxic effect on tumor-bearing mice. Food Chem Toxicol 50: 3166–3173

